# Protein sequence optimization with a pairwise decomposable penalty for buried unsatisfied hydrogen bonds

**DOI:** 10.1101/2020.06.17.156646

**Authors:** Brian Coventry, David Baker

## Abstract

In aqueous solution, polar groups make hydrogen bonds with water, and hence burial of such groups in the interior of a protein is unfavorable unless the loss of hydrogen bonds with water is compensated by formation of new ones with other protein groups. Hence, buried “unsatisfied” polar groups making no hydrogen bonds are very rare in proteins. Efficiently representing the energetic cost of unsatisfied hydrogen bonds with a pairwise-decomposable energy term during protein design is challenging since whether or not a group is satisfied depends on all of its neighbors. Here we describe a method for assigning a pairwise-decomposable energy to sidechain rotamers such that following combinatorial sidechain packing, buried unsaturated polar atoms are penalized. The penalty can be any quadratic function of the number of unsatisfied polar groups, and can be computed very rapidly. We show that inclusion of this term in Rosetta sidechain packing calculations substantially reduces the number of buried unsatisfied polar groups.

## Introduction

Polar groups on the surface of proteins in aqueous solution make favorable hydrogen bonds with water molecules. If these polar groups become buried, either upon folding or binding to another protein, these hydrogen bonds with water must be broken. The energetic penalty of losing h-bonds with water can be offset if a buried polar atom makes a hydrogen bond to another protein atom. We say that this second polar atom “satisfies” the first polar atom. If when buried, the first atom does not make a hydrogen bond, we call it a “buried unsatisfied”.

Modeling the loss in favorable interactions of buried unsatisfied polar atoms is straightforward with explicit solvent models since upon burial, interactions with explicit water molecules are lost. For protein design and other applications where large scale sampling is required and chemical composition (amino acid identity) is changing, implicit solvation models have considerable advantage over explicit models in computational efficiency. The most computationally efficient implicit solvent models are pairwise additive, but identifying and penalizing buried unsatisfied polar atoms is challenging using such models as burial is a collective property. Instead, most current methods use non-pair additive approaches, often involving solvent accessible surface area calculation. The BuriedUnsatsfiedPolarCalculator in Rosetta for instance first calculates which atoms are inaccessible to solvent, and then determines whether or not they are making a hydrogen bond. These methods work well on a fixed protein, but they are not amenable to the implicit-solvation pairwise-decomposable packer of Rosetta[1] or other rotamer-based packing algorithms.

There have been several attempts to capture the energetic cost of buried unsatisfied polar atom penalty in a pairwise manner. The LK solvation model gives all polar atoms a penalty when another atom enters its implicit sphere of solvation[2]. The LK-Ball solvation model takes the LK solvation model and restricts it to positions most critical for hydrogen bonding[3]. A downside to these pairwise methods is that they are intrinsically additive. Instead of a switching behavior where an atom becomes completely buried and can no longer hydrogen bond with water, the burial is gradual and depends on the local density of nearby atoms. These approaches do not specifically penalize buried unsatisfied polar atoms, but instead attempt to model this effect indirectly through balancing the energies of desolvation and hydrogen bonding.

## Materials and Methods

We describe a method for explicitly penalizing buried unsatisfied polar atoms during sidechain packing. We take advantage of the fact that for rotamer-based side chain packing calculations, while the calculated energies must be pairwise-decomposable, the calculations leading to these energies need not be pairwise-decomposable. The method is applicable to fixed-backbone packing trajectories and requires that all rotamers are available before the Monte Carlo trajectory begins.

### Burial region calculation

To assign the buried region in a sequence-independent way so that it can be determined before amino acid sequence design, the protein is first mutated to poly Leucine with Chi1=240 and Chi2=120. Next, the EDTSurf method[4,5] is used to voxelize cartesian space at 0.5Å resolution, determine the molecular surface with a 2.3Å radius sphere, and label each voxel with its depth below the molecular surface. The burial region is then defined as all voxel XÅ below the molecular surface where X is between 3.5 - 5.5Å depending on user preference. Fig 1B shows an example of a 4.5Å burial region on Ubiquitin[6].

**Fig 1.**
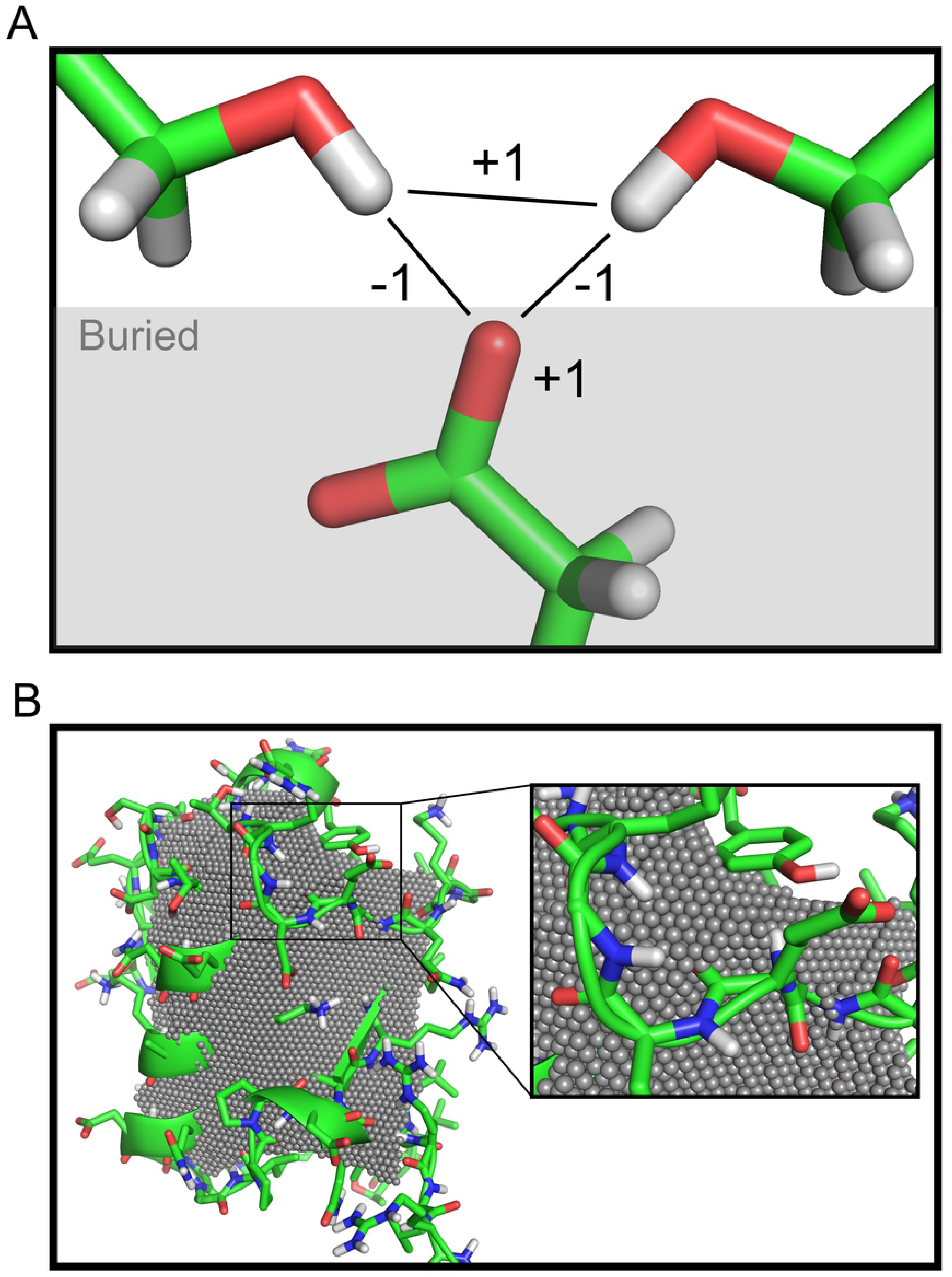
Overview of method. (A) Pictorial representation of the penalty rules. A buried GLU oxygen atom is satisfied by two serine hydroxyls. An oversaturation penalty is applied to the two hydroxyls. In this example: b=1, s=-1, v=1. (B) Burial region calculation applied to Ubiquitin (PDB: 1UBQ)[6]. Grey spheres indicate the buried region at 4.5Å depth from the poly-LEU molecular surface with a 2.3Å probe size.

### Penalty calculation

After the burial region calculation, all buried polar atoms in all rotamers are identified. One-body and two-body atom pseudoenergies can then be assigned with the following simple algorithm:

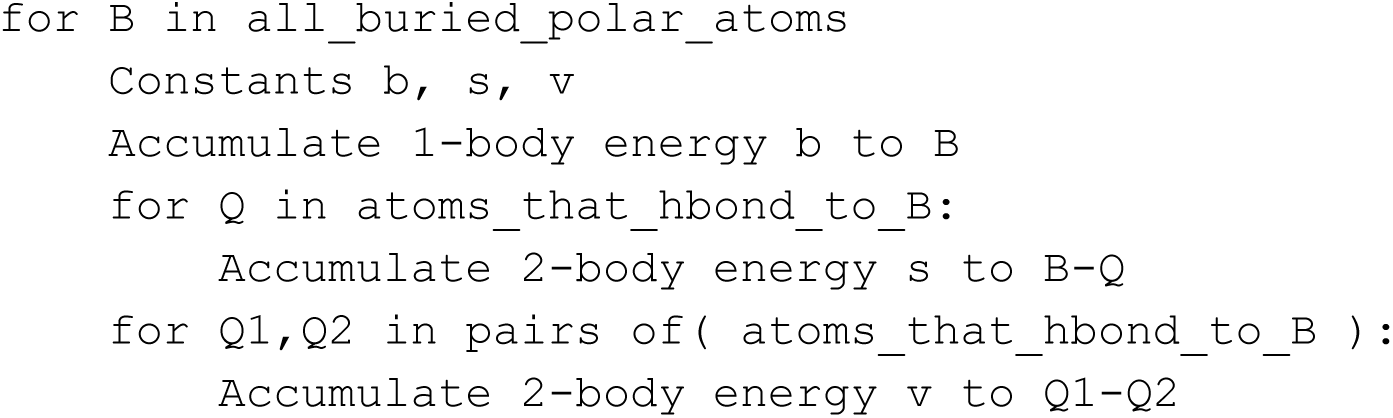

Constants b, s, and v, representing the atom burial penalty, atom-atom satisfaction bonus, and atom-atom oversaturation penalty may be selected on a per-buried-polar-atom basis according to Equation 1. Fig 1A gives a pictorial illustration of this algorithm, which is O(n^3^) on the local number of polar rotamers that h-bond at nearby sequence positions; the algorithm is approximating a 3-body interaction.

### Oversaturation rotamer correction

A problem occurs with this simple algorithm when multiple B from different rotamers at the same sequence position hbond to the same Q1-Q2 pair. With each additional B, the oversaturation penalty between Q1 and Q2 rises. This oversaturation penalty is an error because these B cannot exist at the same time. The solution is to only add the max oversaturation penalty that one rotamer can generate at each position.

#### Corrected Algorithm

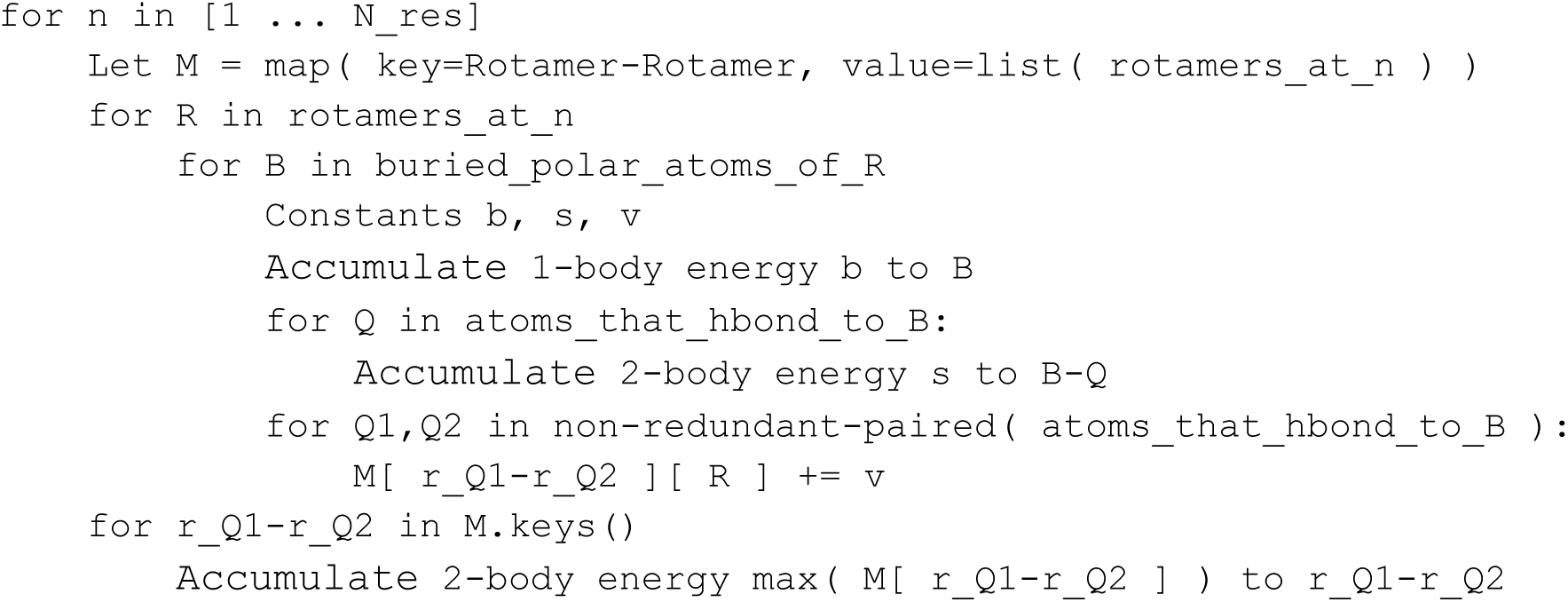

where r_Q refers to the rotamer containing Q. The memory footprint of M can be greatly reduced if instead of storing a list, one only stores the running max value after iterating over each rotamer R.

## Results

### Buried unsatisfied polar quadratic penalty

The algorithm described in the Materials and Methods generates a penalty P of the form

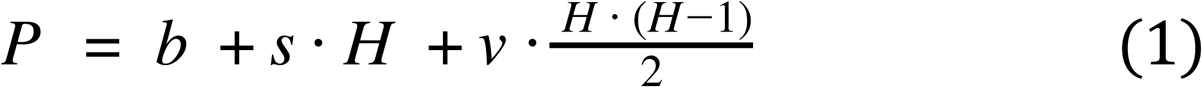

where H is the number of h-bonds, b is the penalty for burying a polar atom, s is the bonus for satisfying a buried polar atom, and v is the penalty for oversaturating a buried polar atom. Because of the quadratic term, this formulation can better describe the “all or none” aspect of buried unsatisfied atoms than linear models such as the LK solvation model. The coefficients b, s, and v can be modified to give any quadratic relationship between the number of h-bonds and the penalty. Additionally, they can be modified on a per-atom basis to give different penalty profiles to different atoms types. As Table 1 shows, parameters can be chosen in general to favor any single number or pair of consecutive numbers of h-bonds.

**Table 1.** Example penalty schemes for different atom types

| Atom Type | Target #H-Bonds | Coefficients | Resulting Penalty for Buried Polar Atom |  |  |  |  |  |
| --- | --- | --- | --- | --- | --- | --- | --- | --- |
| NH1 | 1 | $b=1, s=-1, v=2$ | 1 | 0 | 1 | 4 | 9 | 16 |
| Carbonyl O | 1 or 2 | $b=1, s=-1, v=1$ | 1 | 0 | 0 | 1 | 3 | 6 |
| NH2 | 2 | $b=4, s=-3, v=2$ | 4 | 1 | 0 | 1 | 4 | 9 |
| Carboxylate O | 2 or 3 | $b=3, s=-2, v=1$ | 3 | 1 | 0 | 0 | 1 | 3 |
| NH3 | 3 | $b=9, s=-5, v=2$ | 9 | 4 | 1 | 0 | 1 | 4 |
|  |  | # h-bonds | 0 | 1 | 2 | 3 | 4 | 5 |

Examples of coefficients that lead to a penalty of 1 for buried polar atoms that are off-by-1 from their ideal number of h-bonds. The target # Hbonds listed here are for example only and are not necessarily ideal.

Since the atomic depth calculation is performed on a poly-Leucine backbone, there is no dependence upon the sequence or sidechain conformations to determine burial; hence atomic burial can be pre-computed once before packing begins. However, polar atoms just below the surface will not be considered buried. Consider for instance a backbone carbonyl oxygen that is outside of the burial region, but that is covered by a Phenylalanine ring. Such an oxygen would be buried by explicit solvent or SASA-based measure, but would not be buried by this algorithm.

### Fewer buried unsatisfied polar atoms

One hundred native proteins between 90 and 100 amino acids were redesigned to only allow polar residues as described in S1 File. Fig 2A shows that redesigning the protein with ref2015[7] + the algorithm gave fewer buried unsatisfied polar atoms than designing with ref2015 alone. The 50th percentile structure coming from the algorithm had 2 buried unsatisfied polar atoms while the 50th percentile structure from ref2015 alone had 5. Remarkably, as Fig 2B shows, this reduction from 5 to 2 unsatisfied atoms was accomplished with only 1 to 2 extra h-bonds.

**Fig 2.**
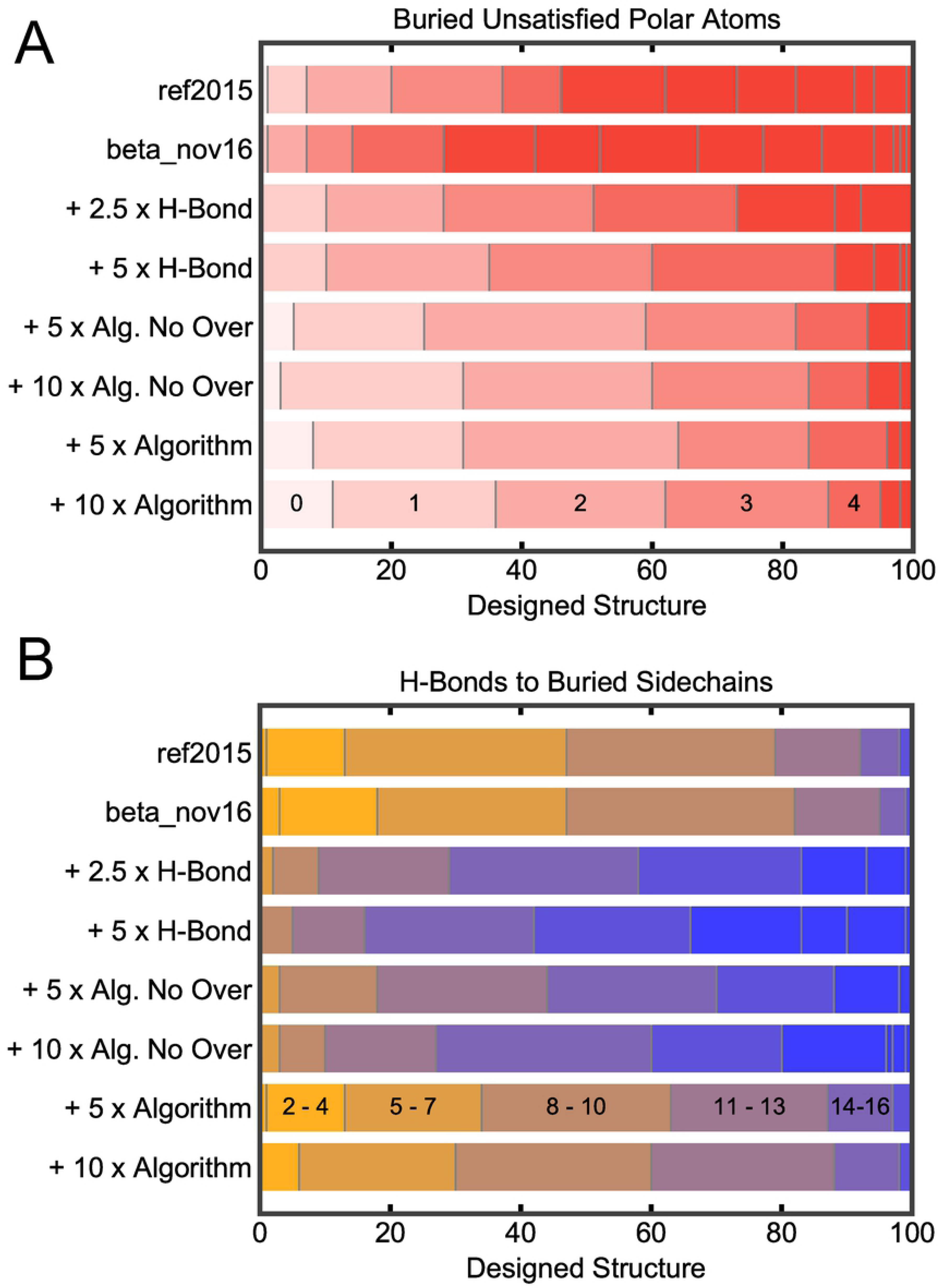
Effect of penalty on native protein redesign. 100 native proteins were redesigned using the energy function and protocol described at the left of the panels, allowing only polar sidechains. All of the methods that start with “+” use ref2015. A) Number of buried unsatisfied polar atoms for each protein. In order from the left, vertical divisions indicate the number of proteins that have 0, 1, 2, or more unsatisfied polar atoms as indicated in the last row. B) Number of h-bonds to buried sidechains. In order from left, each division represents the number of proteins that had from X to (X+2) h-bonds to buried sidechains with each division to the right representing from (X+3) to (X+5). For more information, see S1 File.

### Algorithm is better than simply upweighting h-bonds

A possible objection to the algorithm is that it is simply overcounting h-bonds. This objection was put to the test and Fig 2A shows that the algorithm results in fewer buried unsatisfied polar atoms than simply increasing the hydrogen bond strength. The effectiveness of the oversaturation penalty was also investigated and Fig 2A shows that the algorithm gave roughly the same number of buried unsatisfied polar atoms with and without the oversaturation penalty.

Examining the number of buried h-bonds in Fig 2B; however, shows the effectiveness of the oversaturation penalty. Without the oversaturation penalty, the algorithm makes almost as many h-bonds as simply upweighting h-bonds. After adding the oversaturation penalty, the number of buried hbonds decreases dramatically from a median of 14 to a median of 9 while the number of buried unsatisfied polar atoms remains constant. However, a limitation of our approximation is that the oversaturation penalties do not depend on the presence of the buried polar atom rotamer. For instance, in Fig 1, if the glutamate rotamer was not present, the penalty between serine rotamers would still be applied. To investigate the extent to which this happens, Outer Membrane Phospholipase A of E. Coli, a natural protein with extensive buried polar networks was repacked with an implementation of this algorithm in Rosetta. As Fig 3A shows, the less strict the threshold for h-bonds, the more the extraneous penalties. As the number of rotamers increase, either by adding extra rotamers, or enabling design, the number of extraneous penalties further increases. This problem may be reduced by increasing the stringency of the h-bond-quality threshold or limiting the number of polar rotamers.

**Fig 3.**
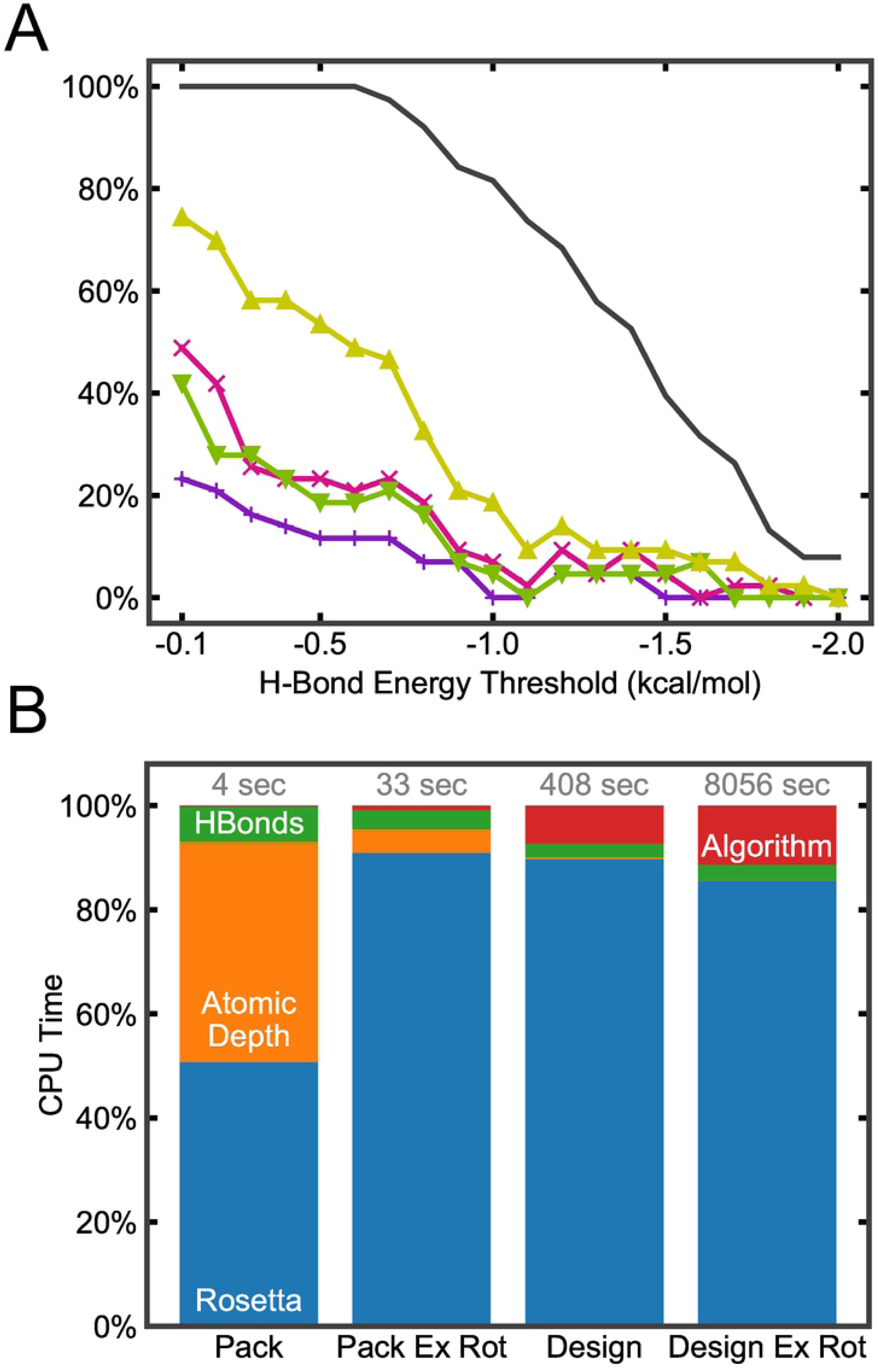
Extraneous oversaturation and performance. A) The Outer Membrane Phospholipase A (PDB: 1ILZ)[8] was either repacked with standard rotamers (purple plus) or extra rotamers (pink cross) or redesigned with all amino acids using standard rotamers (green down arrow) or extra rotamers (yellow up arrow). An expansive buried h-bond network exists in the structure. The percentage of native rotamers in this h-bond network that experience extraneous oversaturation penalties to other native rotamers is plotted vs the energy threshold for a h-bond to be considered. In short, the extraneous oversaturation penalties were determined by performing the algorithm and looking for penalties between native rotamers that were not present before the design/repack rotamers were considered (see S1 File). The black line shows the percentage of h-bonds in the h-bond network that pass the energy threshold. B) Performance of Rosetta-implementation of algorithm. Each stacked bar graph represents the CPU time spent performing a packing or design calculation on 1ILZ. The red top bar represents time spent applying the penalty rules, green bar represents time spent calculating h-bonds between rotamers inside the algorithm, the orange bar is time spent calculating atomic depth, and the blue bottom bar is runtime of the background packing or design calculation. With better data management, the green bar could be avoided as h-bonds are double calculated here (with the other calculation occuring inside blue).

While the extraneous oversaturation penalties may pose a problem for structure prediction, their effect on protein design is not as severe. For designed proteins, there may be several solutions that satisfy a backbone and if one of them is eliminated by an error, there are still several other equally valid solutions, and an almost infinite number of backbones may be attempted. Even if all solutions for a backbone are eliminated another backbone will likely provide solutions. Overall, for design, sins of omission (moving forward with designs containing buried unsats) are more serious than sins of commission (incorrectly eliminating a reasonable design).

### Runtime cost is small

Fig 3B shows the runtime-performance of an implementation of this algorithm as compared to the ref2015 energy function in Rosetta for packing. For very short packing trajectories, the atomic depth calculation dominates the runtime. However, since the atomic depth calculation takes a constant 1.5 seconds on this protein, once the number of rotamers grows, atomic depth is no longer a significant factor. The O(n^3^) scaling of the algorithm, compared to the normal O(n^2^) scaling of pairwise terms, becomes apparent here as the percentage of the runtime increases with increasing rotamers. Even still, the algorithm only consumes 10% of the runtime for the moderate sized case and 16% of the runtime for the extreme case of allowing all amino acid types at every position with extra rotamers.

## Discussion

An advantage of an explicit penalty for buried unsatisfied polar atoms is that protein designers can set the penalty to values appropriate to their application. While many of the previous approaches sought to correctly model the energetics of protein electrostatics, their final functional form was intertwined with the rest of the force-field in a way that could not be arbitrarily modified. Explicit control allows designers to eliminate buried unsatisfied polar atoms independent of overall protein energetics. Designers also have control over the level of hydrogen bond satisfaction in their designs. For example, the number of h-bonds that the NH2 group of glutamine (1, 2, or 1 or 2) must make to be considered satisfied can be specified by suitable parameter choices (Table I). While this paper and the implementation in Rosetta do not consider explicit bound water molecules in crystal structures or from other sources, incorporating these is very simple. As long as the location of the water molecules is known at pre-compute time, they may simply be modeled as polar atoms that can make hydrogen bonds. In its current form without explicit water atoms, the algorithm is now widely used in our research group, and we expect that it will quite broadly help address long standing issues with buried unsaturated polar atoms in de novo protein design.

## Acknowledgements

We thank Vikram K. Mulligan and Scott Boyken for helpful thoughts and conversations about buried unsaturated polar atoms and the idea of creating an energetic penalty for these.

## Supplemental Information

**S1 File. Data collection for figures**

**S2 File. Scripts to produce results**

**S3 File. Results data**

